# The relationship between affective visual mismatch negativity and interpersonal difficulties across autism and schizotypal traits

**DOI:** 10.1101/2021.02.11.430861

**Authors:** Talitha C. Ford, Laila E. Hugrass, Bradley N. Jack

**Affiliations:** Cognitive Neuroscience Unit, School of Psychology, Deakin University, Geelong, Victoria, Australia; Centre for Human Psychopharmacology, Faculty of Heath, Arts and Design, Swinburne University of Technology, Melbourne, Victoria, Australia; Department of Psychology and Counselling, School of Psychology and Public Health, La Trobe University, Melbourne, VIC, Australia; Research School of Psychology, The Australian National University, Canberra, Australia

**Keywords:** visual mismatch negativity, electroencephalography, autism, schizotypy, facial emotion processing

## Abstract

**Objective:** Sensory deficits are a feature of autism and schizophrenia, as well as the upper end of their non-clinical spectra. The mismatch negativity (MMN), an index of pre-attentive auditory processing, is particularly sensitive in detecting such deficits; however, little is known about the relationship between the visual MMN (vMMN) to emotional faces and autism and schizophrenia spectrum symptom domains.

**Methods:** We probed the vMMN to happy, sad, and neutral faces in 77 young adults (18-40 years, 41 female), and evaluated their degree of autism and schizophrenia spectrum traits using the Autism Spectrum Quotient (AQ) and Schizotypal Personality Questionnaire (SPQ). Results: The vMMN to happy faces was significantly larger than the vMMNs to sad and neutral faces (*p* > 0.05). The vMMN to happy faces was associated with interpersonal difficulties as indexed by AQ Communication and Attention to Detail subscales, and SPQ Interpersonal Features (*p* < 0.05, *uncorrected*), such that a larger vMMN was associated with more interpersonal difficulties.

**Conclusions:** Pre-attentive processing of positive affect might be more specific to the interpersonal features associated with autism and schizophrenia.

**Significance:** These findings add valuable insights into the growing body of literature investigating symptom-specific neurobiological markers of autism and schizophrenia spectrum conditions.

**Highlights:**

- The visual Mismatch Negativity (vMMN) is larger in response to happy compared to sad or neutral facial expressions
- The vMMN to happy facial expressions is specifically associated with more social communication difficulties
- vMMN to positive affect may be a neural mechanism associated with social communication difficulties in autism and schizophrenia

---

Several lines of evidence point to poor sensory processing in autism and schizophrenia spectrum disorders, particularly pre-attentive processing of visual and auditory information (Jahshan et al., 2012; Lanillos et al., 2020; Randeniya et al., 2018; Thye et al., 2018). The mismatch negativity (MMN) is a well-established marker of pre-attentive processing of auditory information. To elicit a MMN, typically the repeated presentation of a predictable “standard” stimulus is interrupted at random intervals by an unpredictable “deviant” stimulus, and the MMN represents the difference in the neural response to the standard and deviant stimuli (Naatanen et al., 2007, 2012). Several theories attempt to explain the MMN, such as neural fatigue, neural population sharpening, facilitation of stimulus processing, and cortical predictive coding (see Kremláček et al., 2016; Naatanen et al., 2007; Stefanics et al., 2014). The most popular explanation for the MMN is the predictive coding theory, which proposes that the brain generates a probabilistic model of the sensory input (i.e., the standard stimuli), which explains the attenuation of the neural response to the standard over time. The deviant stimulus is a violation of the probabilistic model, and hence results in a larger neural response (Friston, 2005).

Deficits in the auditory MMN have been consistently replicated across the schizophrenia spectrum (Naatanen et al., 2011; Umbricht & Krljes, 2005) – from chronic illness to high-risk groups (Fulham et al., 2014; Jahshan et al., 2012; Lavoie et al., 2018). Deficits in the auditory MMN in autism are less well established, however, with inconsistencies between studies attributed to the heterogeneity of samples investigated (Schwartz et al., 2018). Nevertheless, recent meta-analyses reported generally reduced auditory MMN in children with ASD, but not adults (Chen et al., 2020; Schwartz et al., 2018). Importantly, the auditory MMN has been associated with poorer social/interpersonal functioning in both autism (Fan & Cheng, 2014) and schizophrenia (Featherstone et al., 2017; Lee et al., 2014), as well as in non-clinical groups (Ford et al., 2017).

Far fewer studies have investigated whether the visual counterpart of the auditory MMN, the vMMN, is similarly affected in autism and schizophrenia. In schizophrenia studies, a smaller vMMN has been reported in response to changes on motion direction (Urban et al., 2008), letters (Neuhaus et al., 2013), happy and fearful facial expressions (Csukly et al., 2013; Vogel et al., 2018), and sequence of stimulus presentation (Vogel et al., 2018), but not to changes in Gabor patch orientation (Farkas et al., 2015). The vMMN amplitude to motion direction has also been associated with negative symptom severity (Urban et al., 2008), and the vMMN to happy deviants was associated with better facial emotion recognition (Csukly et al., 2013).

Studies investigating the vMMN in autism have reported reduced vMMN amplitude in adults (Cléry, Roux, et al., 2013) and longer vMMN latency to object shape deformation in children with ASD (Cléry, Bonnet-Brilhault, et al., 2013). No difference in vMMN amplitude or latency was reported for windmill pattern deviants in a small sample of adults (n = 11) with ASD (Maekawa et al., 2011). One study investigating the relationship between facial emotion vMMN and autistic traits reported a reduced vMMN to happy but not sad deviants that was associated with higher overall levels of autistic traits, as quantified using the autism spectrum quotient (AQ) (Gayle et al., 2012). These findings, along with those demonstrating that the vMMN to happy deviants is associated with better facial emotion recognition (Csukly et al., 2013), suggest that the vMMN elicited by changes in emotional expression might index affective reactivity and social competency (Gayle et al., 2012).

Although there is some evidence to suggest that pre-attentive facial emotion processing is associated with autism and schizophrenia spectrum symptomatology, the specificity of the vMMN differences to autism and schizophrenia spectrum symptom dimensions remains unknown. This study, therefore, extends that of Gayle et al. (2012) by investigating the extent to which pre-attentive processing of changes in facial emotions are associated with specific autism and schizophrenia spectrum traits. It was predicted that a smaller vMMN to happy face deviants would be associated with higher autism and schizophrenia trait severity, particularly in the social domain. The extent to which vMMN to happy and sad face deviants predict the positive and disorganised schizotypy dimensions and attention and imagination dimensions of autism were explored.

## Methods

Ethical approval for this study was granted by the Swinburne University Human Research Ethics Committee (2016/092) in accordance with the Declaration of Helsinki. All participants provided written informed consent prior to commencing the study.

### Participants

A total of 77 (41 female, 36 male) health adults aged 18-40 years participated in this study (age mean (*SE*): total = 24.96 (0.58), male = 25.39 (0.86), female = 25.59 (0.78)). All participants had normal or corrected to normal vision, and were free from psychotropic medications and psychiatric illness, except for one male participant taking an SSRI for mild depression who was excluded from analyses. One male participant was removed due to incomplete psychometric data, and an additional 15 were excluded due to poor EEG data (9 female, 6 male). See Table 1 for final sample participant characteristics, including Mann-Whitney *U* non-parametric tests to identify sex differences in continuous variable, and *χ*^2^ to identify sex differences in handedness. Males scored slightly higher than females on AQ Attention Switching and SPQ Disorganisation dimensions (*p* < 0.05).

**Table 1.** Final sample characteristics for full-scale AQ and SPQ total and dimension scores.

|  | Total<br>Mean(*SE*) | Female<br>Mean(*SE*) | Male<br>Mean(*SE*) | Mann-Whitney *U* | *p*-value |
| --- | --- | --- | --- | --- | --- |
| N | 60 | 32 | 28 |  |  |
| Age | 25.02(0.6) | 24.91(0.84) | 25.14(0.86) | 435.0 | 0.853 |
| AQ total | 105.28(2.93) | 99.03(3.84) | 112.43(4.16) | 306.5 | 0.037* |
| Social Skills | 20.2(0.81) | 19.16(1.04) | 21.39(1.25) | 350.0 | 0.148 |
| Communication | 19.13(0.77) | 17.69(0.99) | 20.79(1.15) | 321.5 | 0.061 |
| Attention Switching | 23.28(0.67) | 21.34(0.91) | 25.5(0.81) | 240.0 | 0.002* |
| Attention to Detail | 23.2(0.72) | 22.41(1.06) | 24.11(0.96) | 358.5 | 0.186 |
| Imagination | 19.47(0.68) | 18.44(0.75) | 20.64(1.14) | 375.0 | 0.281 |
| SPQ total | 145.85(4.94) | 136.75(6.29) | 156.25(7.4) | 323.5 | 0.066 |
| Cognitive-Perceptual | 56.82(2.4) | 52.88(2.99) | 61.32(3.72) | 340.0 | 0.111 |
| Interpersonal | 54.63(1.92) | 52.34(2.39) | 57.25(3.05) | 380.5 | 0.320 |
| Disorganisation | 34.4(1.4) | 31.53(1.68) | 37.68(2.18) | 315.0 | 0.049* |
| Handedness (R:L) | 53:7 | 26:6 | 27:1 | $\chi^2 = 3.34$ | 0.068 |

Notes: *SE* = standard error, AQ = autism spectrum quotient, SPQ = schizotypal personality questionnaire, $\chi^2$ = chi square test.

### Measures

The full 50-item autism spectrum quotient (AQ) was used to measure the five autism trait dimensions of Social Skill, Communication, Attention Switching, Attention to Detail and Imagination (Baron-Cohen et al., 2001). The 74-item schizotypal personality questionnaire (SPQ) was used to measure schizophrenia domains of Cognitive-Perceptual Features, Interpersonal Features and Disorganised Features (Raine, 1991). AQ and SPQ items were pseudo-randomised and presented on a 4-point Likert scale from 1 (*strongly disagree*) to 4 (*strongly agree*) in order to improve reliability (Ford & Crewther, 2014; Wuthrich & Bates, 2005). Items were scaled such that *strongly agree* and *agree* = 1, and *strongly disagree* and *disagree* = 0. All statistical analyses were replicated using the full scale scoring system, which retained 1-4 scores for each item, and is presented in the supplementary material. Participants trait depression, anxiety and stress scores were collected using the DASS-21 (Lovibond & Lovibond, 1995), in order to control for these effects on the relationships between AQ and SPQ domains and the vMMN.

### Stimuli

Face stimuli were selected from the Nimstim database of standardised facial expressions (Tottenham et al., 2009). Five grey-scale female faces were selected; we used female faces only to remove gender bias toward faces. All faces were of caucasian appearance, and the happy, sad and neutral emotions for each face was included. We used the SHINE toolbox for Matlab to match faces for mean luminance (Willenbockel et al., 2010).

### EEG procedure

Participants were seated 57 cm from the stimulus presentation computer screen (inside an electrically shielded room). Participants were instructed to focus on a fixation cross (i.e., “+”) in the centre of the screen that randomly alternated in colour between red and green, and to respond as quickly as possible via keypress when the cross changed colour. The face stimuli for the vMMN task were presented directly behind the fixation cross during the colour-change distractor task.

The vMMN task consisted of 6 blocks containing a total of 750 trials (600 standards, 75 deviants, 75 oddballs), with each facial emotion serving as standard, deviant and oddball across the blocks (e.g., neutral standard - happy deviant - sad deviant, neutral standard - sad deviant - happy deviant, happy standard - neutral deviant - sad deviant, etc.). Each face identity was presented an equal number of times in each block (i.e. 120 presentations of each face as standard emotion, and 15 presentations of each face as deviant emotion). Block order was counterbalanced across participants using a randomised balanced Latin-square. Face stimulus presentation was pseudo-randomised, ensuring that the same identity was not shown on consecutive trials, and that standard emotions were presented for the first three and final two trials of each block, and at least two standard emotions were presented between each deviant emotion. Face stimuli were presented for 400 ms, with an inter-stimulus interval of 150 ms and a jitter of ± 50 ms. The participant’s task was to attend to a fixation cross in the centre of the face stimuli, and to press one of two keys when the colour of the fixation cross changed from red to green, or vice versa (see Figure 1). This task ensured that participants were looking at the face stimuli, but were not attending to it. Analyses of the key press data confirmed that the distractor task was performed to a satisfactory level throughout the vMMN recording, with all participants responding within two seconds on at least 80% of trials (M = 97.93%, SD = 2.75%, M_RT_= 650ms SD_RT_ = 99ms).

**Figure 1:**
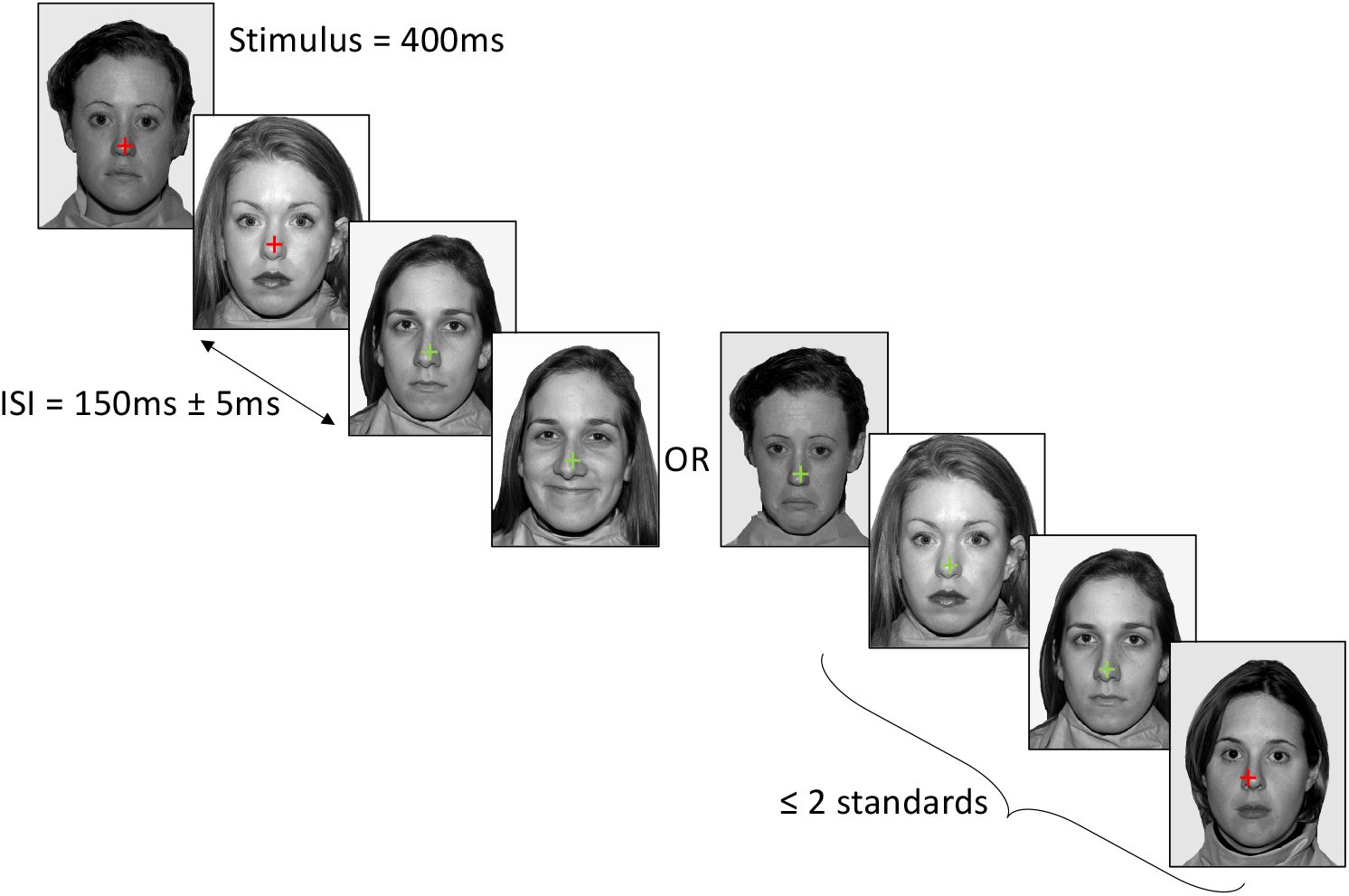
Example stimulus presentation in the neutral standard - happy deviant - sad deviant condition. Stimuli were presented for 400 ms with a 150 ms ± 5 ms jitter. At least 2 standard faces were presented between each deviant. Participants attended to the cross in the centre of the face and pressed the relevant key when the cross changed colour.

### ERP pre-processing

The EEG data were pre-processed in BrainVision Analyzer, Version 2.1 (Brain Products GmBH, Germany). We re-referenced the data offline to the average of all scalp electrodes, and filtered the data using a half-amplitude 0.1 to 30 Hz phase-shift free Butterworth filter (24 dB/Oct slope). EEG data were epoched from -100 to 500 ms in relation to face stimulus onset, and epochs were baseline-corrected to their mean voltage from -100 to 0 ms. Epochs with signals exceeding peak-to-peak amplitudes of 100 microvolts at EEG or EOG channel were excluded. This resulted in the rejection of 14% of all trials. All standard trials immediately following a deviant were excluded from analyses.

ERPs for each emotion (happy, neutral, sad) and for each context (standard, deviant) were computed from occipital electrodes (PO7, O1, Oz, O2, PO8). The time-window for the vMMN was identified using a collapsed localiser approach (Luck & Gaspelin, 2017). All standards and deviants were collapsed across all electrodes and facial emotions, and a difference wave (vMMN) was calculated by subtracting the collapsed standards from the collapsed deviants. The vMMN was then identified as the negative peak between 150 ms and 300 ms, which occurred at 237 ms. The vMMN time-window was then determined as the 100 ms around the vMMN peak; i.e., 187-287 ms. This time-window was used to analyse the uncollapsed vMMNs at each electrode, which were calculated by substracting the mean standard ampitude from the mean deviant amplitude for each facial emotion (i.e., happy vMMN = happy deviant – happy standard).

### Statistical analysis

An *a priori* power analysis conducted using the pwr package in R (Champely, 2018) indicated that 63 participants were required to detect a moderate effect (*r* =.343, based on Gayle et al. [2012]) with 80% power using correlations with alpha = .05.

A 5 (electrode: PO7, O1, Oz, PO8, O2) x 3 (emotion: happy, sad, neutral) linear mixed effects analysis was conducted to investigate vMMN differences across electrodes and emotions, with subject entered as a random factor. To investigate the extent to which the happy, sad, and neutral vMMN at each electrode were related to autism and schizophrenia trait domains, non-parametric Spearman rank order correlations were calculated. We calculated non-parametric correlation coefficients due to significantly skewed distributions of most trait domains and vMMN values. Trait depression, anxiety and stress were not related to the happy or sad vMMN at any electrode site, thus were not added as covariates in the analyses. Family-wise error was corrected for by applying false discovery rate (FDR; Benjamini & Hochberg, 1995). All statistical analyses were conducted using Jamovi version 1.2 (*Jamovi*, 2020).

## Results

The grand mean average vMMN ERP for each facial emotion at each visual electrode are shown in Figure 2, and the electrode x emotion mixed effects plot of mean vMMN is shown in Figure 3. There was a clear negativity between 150 ms and 300 ms for both happy and sad facial expressions, with the happy vMMN appearing to peak earlier than the sad vMMN.

**Figure 2:**
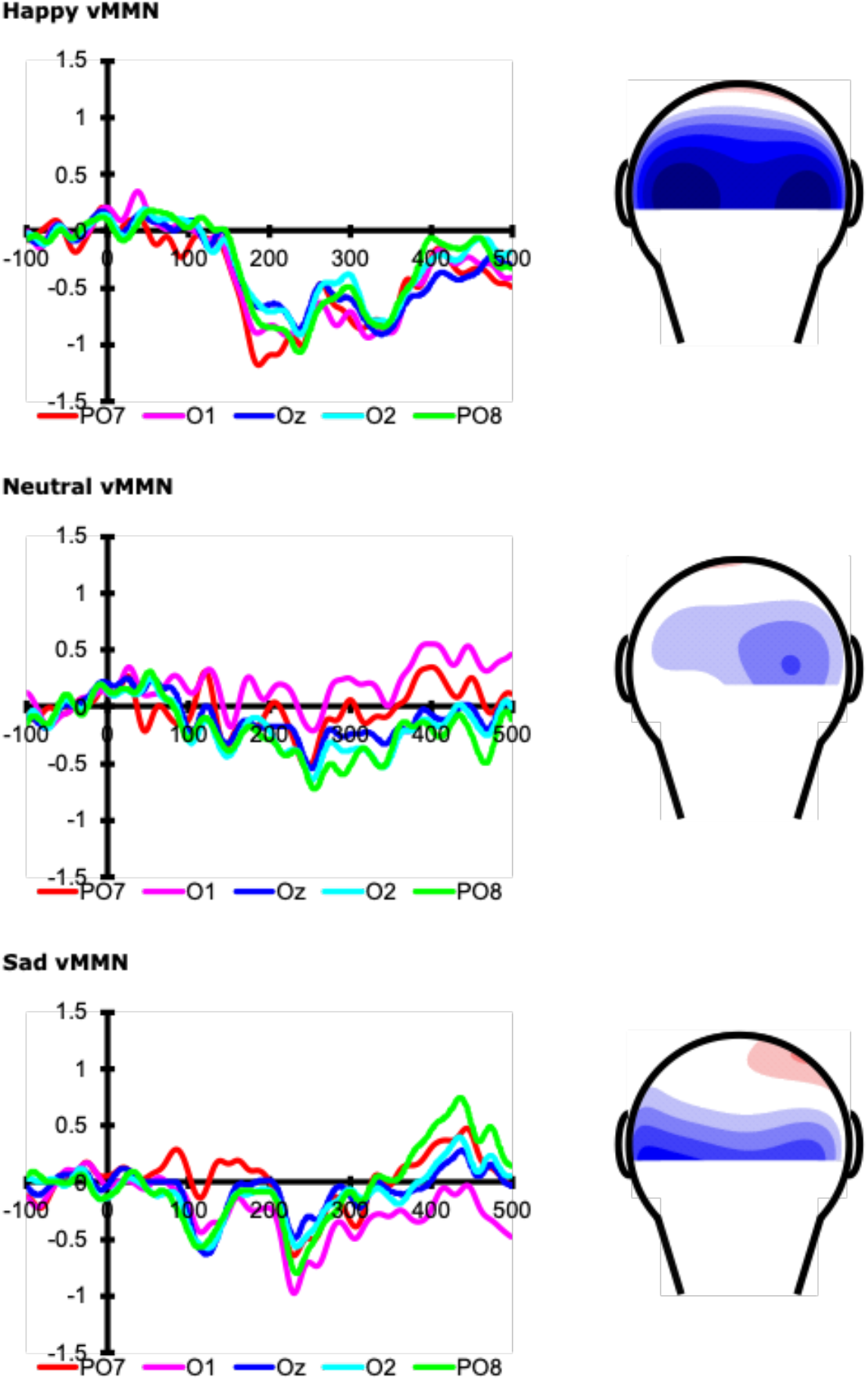
The line graph shows the grand-averaged deviant-minus-standard difference waves at occipital electrodes (PO7, O1, Oz, O2, PO8), showing time (ms) on the x-axis, with 0 indicating stimulus onset of the emotional face, and voltage (μV) on the y-axis, with positive voltages plotted upwards. The voltage maps show the distribution of voltages over the back of the scalp during the vMMN time-window (187 to 287 ms).

**Figure 3:**
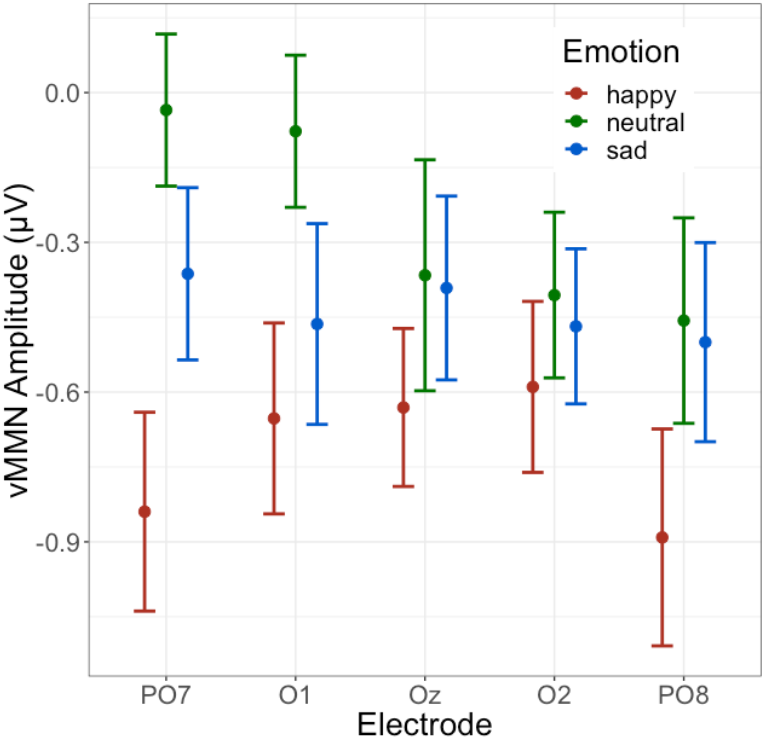
Mean and standard error bars for visual mismatch negativity (vMMN) amplitude (μV) by visual electrode and facial emotion.

The linear mixed effects analysis (electrode x emotion) revealed a significant main effect for emotion (*F*(2, 824.11) = 8.10, *p* < 0.001), with the happy vMMN significantly larger than the neutral vMMN (*t*(824.08) = -3.99, *p* < 0.001) and sad vMMN (*t*(824.08) = -2.48, *p* = 0.013). There was no difference between sad and neutral vMMN (*p* = 0.132). No significant hemisphere main effect or emotion x hemisphere interactions was found (*p*s < 0.1; Figure 3).

Spearman rank order correlations investigating the relationship between the vMMN to happy, sad, and neutral facial expressions found significant moderate negative correlations between the happy vMMN at O1, Oz and O2 and social autism and schizophrenia spectrum domains (*p* < 0.05; see Table 2), indicating that those with more social and communication difficulties and more attention to detail exhibit a larger (more negative) vMMN at O1 and O2 (Figure 4); however, these correlations did not survive correction for FDR. There were no relationships between sad and neutral vMMN amplitude at any electrode site and any of the AQ or SPQ domains (*p* > 0.05).

**Table 2.**
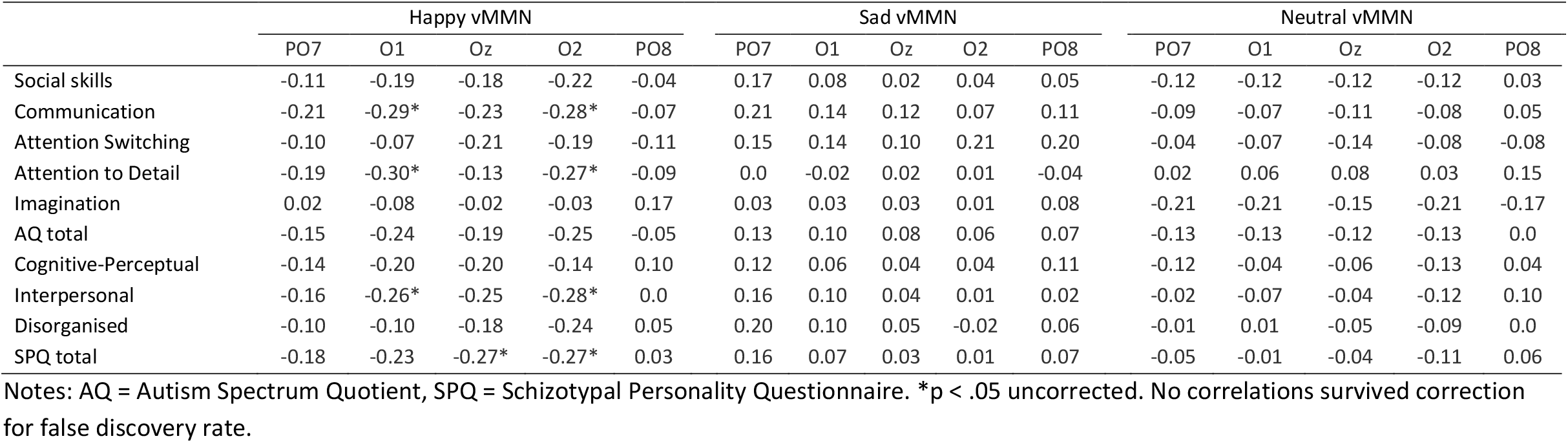
Spearman rank order correlations between AQ and SPQ dimensions and average vMMN amplitudes across visual electrode sites for each emotional expression.

**Figure 4:**
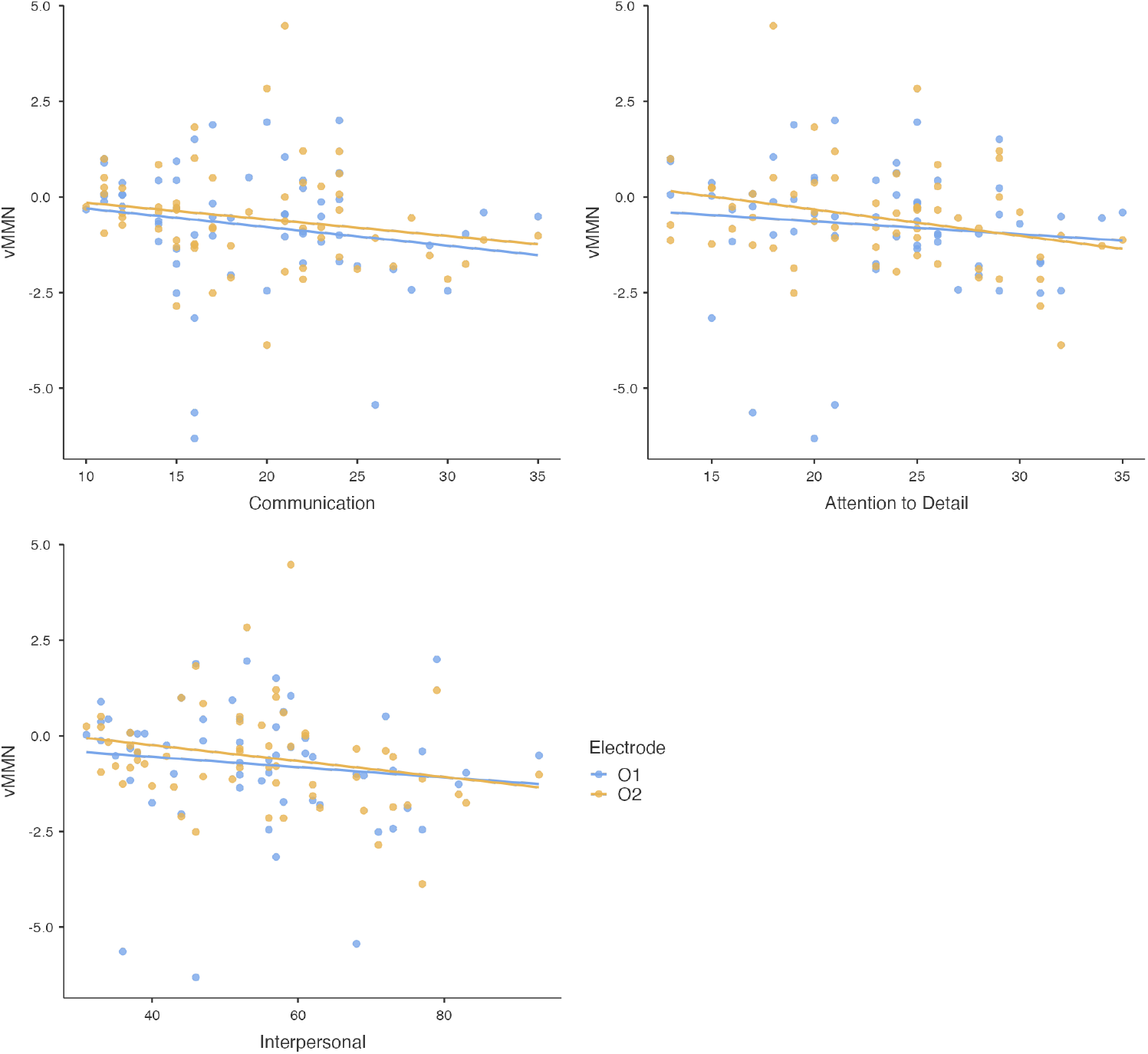
Scatter plots of uncorrected correlations between visual mismatch negativity (vMMN) amplitude (μV) to happy faces and Communication, Attention to Detail and Interpersonal subscales at O1 and O2 electrodes.

In order to compare the current data with previous results (Gayle et al., 2012), we re-ran the AQ and SPQ correlations with the vMMN calculated for happy and sad deviants relative to the neutral standards (Table 3). Similar to the original analyses (i.e., when the vMMN was calculated for the same emotion when it was deviant relative to when it was the standard; see Table 2), the communication domain of AQ and the total SPQ score were negatively correlated with the vMMN at O2 (*p* < .05). There were no other significant correlations.

**Table 3.** Spearman rank order correlations between AQ and SPQ dimensions and average vMMN amplitudes when the mismatch response was calculated for happy and sad deviants relative to neutral standards

|  | Happy vMMN |  |  |  |  | Sad vMMN |  |  |  |  |
| --- | --- | --- | --- | --- | --- | --- | --- | --- | --- | --- |
|  | PO7 | O1 | Oz | O2 | PO8 | PO7 | O1 | Oz | O2 | PO8 |
| Social skills | -0.11 | -0.16 | -0.10 | -0.25 | -0.13 | 0.06 | 0.05 | -0.04 | -0.04 | -0.02 |
| Communication | -0.11 | -0.16 | -0.16 | -0.27* | -0.09 | 0.10 | 0.03 | 0.05 | -0.02 | -0.01 |
| Attention Switching | 0.03 | 0.05 | 0.06 | 0.01 | 0.04 | 0.02 | 0.10 | 0.06 | 0.11 | 0.10 |
| Attention to Detail | -0.09 | -0.13 | -0.14 | -0.15 | -0.02 | -0.04 | 0.01 | 0.09 | -0.02 | -0.11 |
| Imagination | 0.05 | 0.02 | 0.16 | 0.02 | 0.22 | -0.07 | 0.01 | -0.03 | -0.04 | -0.03 |
| AQ total | -0.07 | -0.10 | -0.05 | -0.18 | -0.01 | 0.01 | 0.05 | 0.03 | -0.02 | -0.04 |
| Cognitive-Perceptual | -0.09 | -0.22 | -0.21 | -0.23 | 0.04 | -0.02 | -0.05 | -0.07 | -0.12 | -0.10 |
| Interpersonal | -0.04 | -0.18 | -0.13 | -0.24 | -0.08 | 0.01 | -0.03 | 0.02 | -0.12 | -0.14 |
| Disorganised | 0.10 | -0.01 | -0.09 | -0.24 | 0.07 | 0.03 | -0.04 | -0.03 | -0.12 | -0.16 |
| SPQ total | -0.07 | -0.21 | -0.21 | -0.29* | -0.02 | -0.12 | -0.05 | -0.02 | -0.14 | -0.17 |

Notes: AQ = Autism Spectrum Quotient, SPQ = Schizotypal Personality Questionnaire. \* $p < .05$ uncorrected. No correlations survived correction for false discovery rate.

## Discussion

This study is the first to investigate the relationship between emotion-related vMMN amplitudes and specific autism and schizophrenia spectrum traits within the non-clinical population. This work extends from Gayle et al. (2012) who reported an association between the emotion-related vMMN and total AQ scores only. As expected, there were strong bilateral occipitotemporal vMMN responses to the happy and sad expressions. Although vMMN amplitudes were not significantly correlated with the total AQ score, we found that people with higher levels of interpersonal difficulty specifically (as indexed by the AQ Communication and Attention to Detail, and SPQ Interpersonal Features subscales) tended to exhibit stronger vMMN responses to happy expressions. Despite the large sample size, however, none of these correlations survived correction for family-wise error. Given the theoretical importance of an association between autistic traits and vMMN amplitudes, it is necessary to consider why the correlations between autistic personality traits and the vMMN are in the opposite direction to those previously reported.

These data reveal a weak-moderate relationship between interpersonal difficulties and the vMMN to happy faces in the opposite direction to Gayle et al. (2012). A notable methodological difference between the two studies is that the previous study calculated the vMMN responses to happy and sad deviants relative to neutral standards. Hence, differences in the morphology of the neural response to the standard and deviant might reflect differences in response to the specific stimulus being presented (i.e., a sad face versus a neutral face), as opposed to a neural response to prediction violation. It has been shown that within the vMMN time-window, evoked responses to emotional faces tend to be lower amplitude for people with ASD or high levels of autistic traits than for people with low levels of autistic traits (Csukly et al., 2013; Gayle et al., 2012). Hence, Gayle et al.’s results may have reflected a correlation between AQ and the evoked responses to facial emotional, rather than a correlation between AQ and responses to emotion prediction error. However, our subsequent analyses showed that regardless of whether we compared deviants and standards for the same emotion (i.e., happy deviant vs happy standard), or compared emotional deviants with neutral standards, correlations between the happy vMMN amplitude and autistic and schizotypal personality traits tended to be negative. Thus, our results indicate that predictive error responses to unexpected happy faces tend to be stronger for people with higher levels of interpersonal difficulties, as indexed by the subscales of the AQ and SPQ.

Although the increase in vMMN to happy deviants (compared to happy standards) with more social and interpersonal difficulties was unexpected, previous studies of the auditory vMMN in Asperger’s Syndrome (AS) or ASD have also reported conflicting findings. For instance, Lepisto et al. (2007) found that adults with AS tended to exhibit enhanced MMNs to speech sounds, but not for non-speech sounds, suggesting an auditory hypersensitivity or filtering difficulty in AS (Lepistö et al., 2007). Furthermore, Schwartz and colleagues’ (2018) meta-analysis found that while the auditory MMN was generally reduced in children with ASD, adults with ASD elicit a larger MMN to speech deviants compared to controls. However, the meta-analysis highlighted that many studies of the auditory MMN had small sample sizes and did not properly counterbalance their stimuli.

We observed moderate relationships between autistic and schizotypal traits and emotion-related vMMN amplitudes for happy facial expressions (uncorrected), but not for sad or neutral expressions. This finding is consistent with Gayle et al., who reported that the total AQ score was selectively related to vMMN amplitude in response to happy expressions but are surprising in relation to evidence that people with ASD exhibit more general deficits in affective processing (Eack et al., 2015; Fan & Cheng, 2014; Wright et al., 2008). It could be that different aspects of affective processing have different relationships with autistic traits; for instance, Wilbarger et al. (2009) found that people with ASD exhibit an atypical startle response to positively valanced faces, despite showing typical implicit responses with automatic facial mimicry and overt valence ratings. This indicates a disruption in an early stage of affective processing, such that there is a dissociation between higher-order assignment of valence and basic automatic defensive responses. The authors suggest that this ambiguity may mean that positive stimuli are experienced as unreliable or potentially harmful, which could in turn lead to interpersonal difficulties. The current results are consistent with the idea that people with higher self-report levels of interpersonal difficulty tend to exhibit a heightened automatic alerting response to unexpected emotions.

The vMMN to happy faces over O1 and O2 was also shown to increase with more AQ Attention to Detail (uncorrected), which is somewhat intuitive given better preattentive discrimination between emotions could lend itself to more superior attention to detail in the social environment. Again, better discrimination of happy deviants among neutral and sad standard faces may not necessarily translate to correct interpretation of facial expressions. Finally, SPQ scores were associated with larger vMMN amplitudes for the happy facial expressions (uncorrected). This effect was also specific to occipital sites (Oz and O2) and was driven by weak-moderate relationships across the three schizotypy dimensions (Cognitive-Perceptual, Interpersonal and Disorganised Features). In contrast, the relationship between AQ subscales Imagination and Attention Switching and the happy vMMN were weak across electrode sites, driving down the relationship between AQ total and the vMMN.

Our design allowed the vMMN to be computed as the difference between deviant and standard of each emotion and removed a particular potential bias toward happy and sad emotion compared to the neutral emotion. A further strength was the use of grey-scale faces of four different female actors. The use of female-only stimuli and equal number of male and female participants recruited in this study removed any potential sex vMMN (Kecskes-Kovacs et al., 2013), and allowed us to be more confident that the vMMN was due to emotion prediction error, rather than a sex prediction error. Furthermore, through investigating the relationship between the vMMN to facial emotions and varying degrees of specific autism and schizophrenia spectrum symptom traits in a non-clinical sample, these findings highlight the utility of this population to better understand the neurobiology of symptoms at a clinical level. Indeed, the relationship between sub-clinical and clinical auditory MMN profiles has been well established, with high-risk individuals and first-degree relatives showing a similar MMN deficit, albeit less pronounced, to clinical groups (Brockhaus-Dumke et al., 2005; Lavoie et al., 2018).

Despite the robust research methods and large sample size employed herein, there are some limitations that should be taken into consideration when interpreting these data. First, we did not include a non-emotional deviant, such as stimulus tint. Instead, we calculated the vMMN by subtracting the deviant of each emotion from the standard of the same emotion (i.e., happy deviant minus happy standard), which ensured the detected vMMN was not simply a response to the physical characteristics of faces or to the change in emotional expression. Furthermore, although participants were excluded if they were taking psychoactive medications, they were not asked to abstain from caffeine, nicotine, alcohol, cannabis, or other illicit substances prior to testing. Only moderate alcohol consumption (.04-.07% BAC) has been shown to affect the vMMN (Kenemans et al., 2010), but acute doses of nicotine, alcohol and cannabis affect the auditory MMN, an thus may also effect the visual counterpart. Given testing for this study occurred on weekdays, it is likely that participants were only acutely affected by caffeine, which has been shown not to affect auditory MMN (Rosburg et al., 2004). Nevertheless, these data should be interpreted with the above limitations in mind. Finally, due to the large number of statistical tests performed, none of the correlations survived correction for FDR. Future research should be conducted on a larger sample, or with more targeted hypotheses, to replicate these findings.

This study sought to further explore the findings of Gayle et al. (2012) by investigating the relationship between the vMMN to happy and sad facial expressions and specific autism and schizophrenia spectrum traits in a large sample of healthy adults. We addressed the research design limitations outlined in previous studies and found that communication and interpersonal difficulties, as well as attention to detail, were associated with a greater vMMN amplitude in response to happy faces, however, the statistical significance of these effects diminished following correction for family-wise error. The large early visual response to happy faces might suggest a hyper-responsivity to the change in emotional expression, with this hyper-responsivity possibly leading to poorer interpersonal outcomes. In utilising a non-clinical population, these findings add valuable insights into the growing body of literature investigating symptom-specific neurobiological markers of autism and schizophrenia spectrum conditions.

## Acknowledgements

We thank Swinburne Neuroimaging for use of the EEG facilities, and our third year Psychophysics students, Tomas Cox, Edenn Baczyk, Jodeci Cowell and Alexandra Suvorova, for their help with recruitment and data collection. Dr Ford is supported by a Deakin University Dean’s Postdoctoral Research Fellowship.

## Declaration of interests

None

## Notes

### Competing Interest Statement

The authors have declared no competing interest.

### Summary of Updates

Following peer review, we have clarified the methods and results sections, and modified the discussion.

